# The Evolution of Antibiotic Production Rate in a Spatial Model of Bacterial Competition

**DOI:** 10.1101/342600

**Authors:** Jakub Kosakowski, Prateek Verma, Supratim Sengupta, Paul G Higgs

## Abstract

We consider competition between antibiotic producing bacteria, non-producers (or cheaters), and sensitive cells in a two-dimensional lattice model. Previous work has shown that these three cell types can survive in spatial models due to the presence of spatial patterns, whereas coexistence is not possible in a well-mixed system. We extend this to consider the evolution of the antibiotic production rate, assuming that the cost of antibiotic production leads to a reduction in growth rate of the producers. We find that coexistence occurs for an intermediate range of antibiotic production rate. If production rate is too high or too low, only sensitive cells survive. When evolution of production rate is allowed, a mixture of cell types arises in which there is a dominant producer strain that produces sufficient to limit the growth of sensitive cells and which is able to withstand the presence of cheaters in its own species. The mixture includes a range of low-rate producers and non-producers, none of which could survive without the presence of the dominant producer strain. We also consider the case of evolution of antibiotic resistance within the sensitive species. In order for the resistant cells to survive, they must grow faster than both the non-producers and the producers. However, if the resistant cells grow too rapidly, the producing species is eliminated, after which the resistance mutation is no longer useful, and sensitive cells take over the system. We show that there is a range of growth rates of the resistant cells where the two species coexist, and where the production mechanism is maintained as a polymorphism in the producing species and the resistance mechanism is maintained as a polymorphism in the sensitive species.

**Author Summary:** Natural environments such as the soil contain many species of antibiotic producing bacteria. Antibiotics prevent the growth of sensitive species that would otherwise outcompete the more-slowly-growing antibiotic producers. The producers are also vulnerable to competition from non-producing “cheats” arising by mutations within the producing species that avoid the metabolic cost of antibiotic production. We consider multiple strains of producers that differ in production rate in the presence of sensitive cells of a different species. We show, in 2d simulations, that the system evolves towards a state with a dominant producer strain that is able to outcompete the sensitive cells, plus a range of low-rate producers and non-producers that can survive in the presence of the dominant producer, but not on their own. This system remains stable, despite the short-term selective advantage to reducing production rate. When resistant mutants are added to the sensitive species, we show that there is a range of growth rate of the resistant cells in which producers, non-producers, sensitive and resistant cells can all coexist - as we see in nature. Our model shows the balance of factors required to maintain resistance mechanisms and production mechanisms together within the mixture of species.

## Introduction

Microbial communities are characterized by remarkable levels of species diversity suggesting a complex interplay between environmental conditions, nutrient sources, community structure and nature of interactions between species making up the community. The diversity of these systems has been widely studied. One study found that 1g of deciduous forest soil contains ~1.5 x 10^10^ bacteria spread across ~4,000 different species [1]. Biogeographical analysis indicates communities vary widely based on geographical location and factors such as pH, and points to the existence of a wide range of complex and diverse ecosystems [2]. The maintenance of such bio-diverse microbial eco-systems is a central problem in ecology.

Of special interest are microbial communities containing antibiotic-producing bacteria [3–5]. Several recent studies have focused on identifying the conditions under which such communities involving antibiotic producers (P) can stably co-exist with antibiotic resistant (R) and antibiotic sensitive (S) strains. This question has become particularly relevant in the light of growing antibiotic resistance of human pathogens. Due to the serious global public health consequences of such antibiotic resistance, it is essential to understand how antibiotic resistant strains can survive and thrive in bacterial communities inhabited by multiple bacterial species.

The rock-paper-scissors game, in which there are three species that each outcompete one of the other two, has proved to be a useful framework for investigating the conditions for biodiversity in a variety of ecosystems, including bacteria [6–8], lizards [9], and salmon [10]. The mobility of competing species [11,12] as well as the structure of the underlying populations [13–17] has been shown to be critical factors in the stable coexistence of multiple species.

Chao and Levin [11] demonstrated that in liquid culture, colicin-producing bacteria survive only at high starting densities, whereas in agar cultures they remain competitive even at low starting densities. Wiener [12] also demonstrated the positive effect clustering has on the antibiotic producer growth when competing with a sensitive strain, and the negative effect of higher starting densities of non-producers of antibiotics (cheaters). Kerr et al. [7] showed using colicin producing (P), sensitive (S) and resistant (R) strains that ecological diversity is sustained only if dispersal is local and is no longer observed in a well-mixed population. They used experiments as well as a lattice-based computational model to demonstrate cyclic dominance of P, S, and R strains in the limit when dispersal is restricted to the local neighbourhood.

Reichenbach et al. [18,19] systematically studied the effect of increasing diffusion rate in three-species systems with deterministic, partially stochastic and fully stochastic (agent-based) dynamics. Remarkably, they found that all three systems show cyclic dominance, manifest through formation of spiral waves, for diffusion rates below a critical threshold. Beyond that threshold, stable coexistence is no longer possible. More recently, Kelsic et al. [20] used a 3- species, 3-antibiotic model to revisit the question of multi-species co-existence in systems where resistant strains function by producing an enzyme that degrades the antibiotic. They showed that the presence of degraders allows for maintenance of biodiversity regardless of dispersal rates and that the system is also stable against invasion by cheater mutants.

Most models discussed above are bactericidal in nature implying that the antibiotic increases the death rate of sensitive (S) species. An alternative to such models are bacteriostatic models where the antibiotic reduces the birth rate of the sensitive strain. In this paper, we consider producer cells that produce antibiotic at a rate *a*. The antibiotic reduces the growth rate of sensitive cells in proportion to the local antibiotic concentration (dependent on *a*), but the growth rate of the producers is also reduced in proportion to *a* due to the metabolic cost of production. As a consequence, non-producers (*i.e.* production cheaters that arise by mutations within the producer species) possess a growth advantage relative to the producers. Initially we consider models with three cell types: a producer that produces antibiotic with a specified rate *a*, a non-producer with *a* = 0, and a sensitive cell type of a different species. We show that coexistence of these three types is possible in a spatial model, provided the production rate lies within an intermediate range. If *a* is either too high or too low, only sensitive cells survive.

A central question in this paper is whether the system with three cell types described above is stable to evolution of *a*. From the point of view of the producers, it pays to produce only the minimum amount of antibiotic necessary to prevent being outcompeted by the sensitive cells. However, in the short term, evolution will always select for cells with lower *a*, and hence reduced cost, provided there are some high-rate producers around that can kill off the sensitive cells. The possibility therefore arises that the short-term selection for low *a* may lead to reduction in *a* to the point at which the producers can no longer compete with sensitive cells. We therefore consider a system in which produces of all rates *a* can arise due to mutations in the producer species. Non-producers are the limiting case of *a* = 0. We find that there is a stable state in this case in which a dominant producer with moderately high *a* coexists with a range of low-rate producers and sensitive cells. The presence of evolution of *a* and multiple producer strains does not lead to the loss of stability of the whole system.

We also consider the possibility that a resistance mechanism evolves in the sensitive species. There are now two kinds of cheats in this system. Sensitive cells are cheats relative to resistant cells because they avoid the cost of resistance. Non-producers are cheats relative to producers because they avoid the cost of production. We show that there are conditions in which all four cell types P, N, S and R coexist. The resistance mechanism is maintained as a polymorphism in the sensitive species and the production mechanism is maintained as a polymorphism in the producer species.

In all the cases considered here, coexistence of more than one cell type depends on the spatial structure. Ordinary Differential Equations describing well-mixed systems are qualitatively different from stochastic spatial models, as they predict that only one cell type survives. We will begin this paper by discussing why the well-mixed ODE models do not give coexistence. This also provides a way to summarize the differences between several possible ways of implementing a rock-paper-scissors relationship between cell types. After that, we define the stochastic spatial models that we use here, and present simulation studies of the evolution of antibiotic production and resistance mechanisms in these stochastic spatial models.

## ODE Models for Three Interacting Species

The term rock-paper-scissors is used rather generally to describe three-species models in which each species beats one of the other two and loses to the third. However, there are several non-equivalent ways of defining ordinary differential equation (ODE) models of this type, and we wish to summarize these different ways before introducing the model we study in this paper.

The model of May and Leonard [21] is a rather general way to describe sets of interacting species:

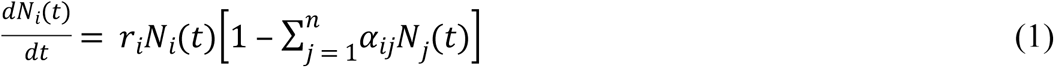

Here, *N_i_*(*t*) is the concentration (number of individuals per unit area) of species *i* at time *t*, *r_i_* is the growth rate of species i, and *α_ij_* is the coefficient describing the effect of species *j* has on species *i*. Several types of behaviour are possible in this model. The outcome that is most relevant here is a heteroclinic cycle that moves around the boundaries of the phase space [22,23]. The system spends most of its time very close to one or other of single-species equilibrium points, and very occasionally switches from one equilibrium to the next. This is only possible because the model is deterministic and allows arbitrarily small densities of each species to recover and become large again. In finite size populations, there can never be fewer than one individual in a population, and once a species is extinct it cannot reappear. Hence, the system is bound to go to one or other of the three single-species equilibrium states.

One well-studied model of this type is that of Reichenbach *et al.* [18,19], for which the ODEs may be written:
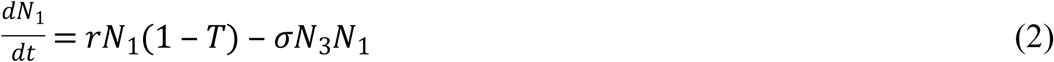

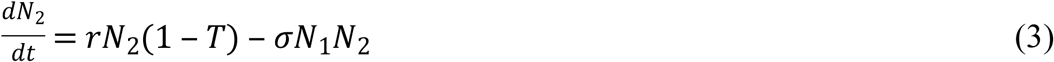

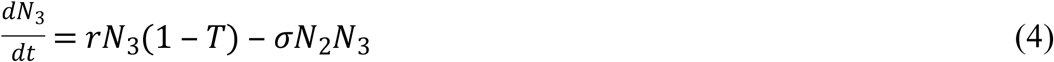

where *T* = *N_1_* + *N_2_* + *N_3_* is the total population density. The terms involving *σ* are further inhibitory interactions in addition to the effects of carrying capacity. The inhibitions occur in a cyclic fashion: 1 inhibits 2, 2 inhibits 3, and 3 inhibits 1. The deterministic infinite system will follow a heteroclinic orbit around the boundaries of the phase space, and a finite size population will tend to one or other of the equilibria with only a single species present. Reichenbach *et al.* [ 18] developed a lattice model whose well-mixed limit is equivalent to the ODEs of equations 2–4, but which shows spiral wave solutions in the spatial case that allow coexistence of the three species.

Another case of three species interactions is the production of colicin [6,7]. In this case the system is composed of three E. coli types: a full producer, *P*, a non-producer (or cheater) *N*, and a sensitive type, *S*. The well-mixed system can be described by a set of ODEs:
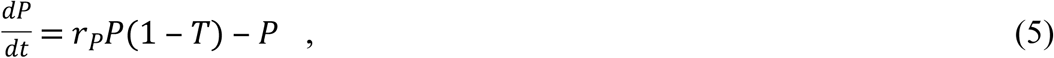

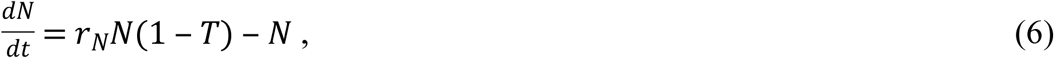

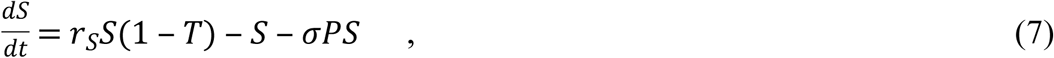

where *T* = *P* + *N* + *S* is the total population density. Here, it is important that *r_P_* < *r_N_* < *r_S_*, and that only the *S* species is inhibited by the colicin production of *P.* Additionally, there is a linear death rate for each species, which is essential because otherwise *T* tends to 1 and no further growth is possible.

If we begin with a mixture of *P* and *S* only, the system in equations 5–7 tends either to a state with only *P* or a state with only *S*, depending on how much *P* is initially present. However, if there is even a small amount of *N* present initially, then *N* always outgrows *P*, so *P* cannot survive. In the absence of *P*, *S* always outgrows *N*. Thus, if we begin from any state with non-zero densities of all three species, the system converges to a steady state with only *S* present, *i.e.* according to well-mixed models, antibiotic producers will always be destroyed by the evolution of non-producing cheaters, which will, in turn, be destroyed by sensitive cells. Durrett and Levin [6] have shown that in a spatial version of the colicin model, it is possible for all three species to coexist for certain ranges of the rate parameters. Patches of the three species move across the lattice and replace one another, but all three remain present in the long term. As with the Reichenbach [18] model discussed above, it is the spatial pattern formation that allows the three species to coexist. However, the patterns of spiral waves and the patterns of random patches that arise in the two cases are qualitatively different.

We now give the ODEs for the well mixed model for the evolution of the antibiotic production rate that we study here.
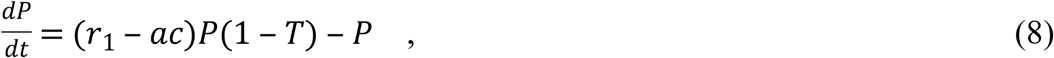

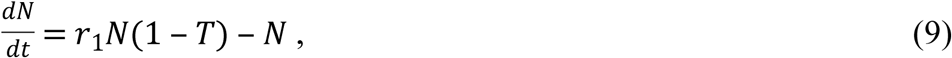

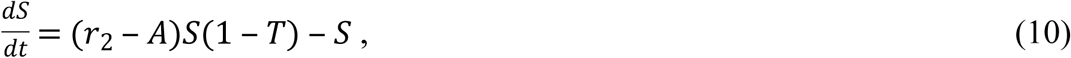

This is similar to the colicin model, but the following points should be noted. Here we are assuming that *P* and *N* are strains of a single species with intrinsic growth rate *r_1_*, whereas *S* corresponds to a second distinct species, with intrinsic growth rate *r_2_* > *r_1_*. The growth rate of *P* is reduced relative to *N* by the cost of antibiotic production *ac*, where *a* is the rate of antibiotic production by *P* cells, and *c* is the metabolic cost per unit of production. We count these two factors separately because we are interested in studying the evolution of the rate of antibiotic production. The parameter *a* is an evolvable property of producers that may be different in different producer strains, whereas *c* is a fixed property of the metabolic pathway for antibiotic production that cannot be changed (unless a completely different antibiotic molecule arises).

The growth rate of sensitive cells in absence of the antibiotic is *r_2_*. When producers are present, this rate is reduced in proportion to the concentration of antibiotic, *A*. Producers produce antibiotic at rate *a*, and the antibiotic breaks down at rate *b*. Hence, 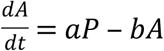. We assume that *a* and *b* are both large relative to the rates of cell division, so that *A* reaches a steady state *A* = *aP/b*. This value of *A* is used in equation 10, which avoids adding a separate ODE for *A*. This is a bacteriostatic case, where antibiotic reduces growth rate of the sensitive cells, rather than a bactericidal case, where the antibiotic increases the death rate (as was the case in equation 7). Note that increasing *a* leads to a decrease in growth rate of both *P* and *S*.

We will now consider the dynamical behaviour of equations 8–10. Firstly, it is clear that there are stable equilibrium solutions for each of the three species separately. If only *P* is present, then *T* = *P*, and the equilibrium density of *P* is:
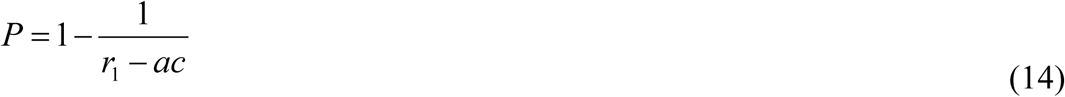

Similarly, the one-species equilibria for *N* and *S* are
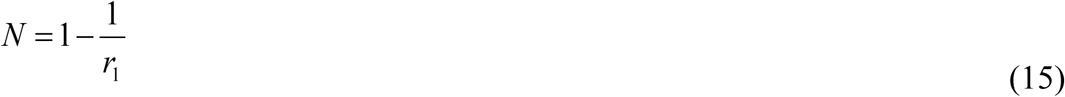

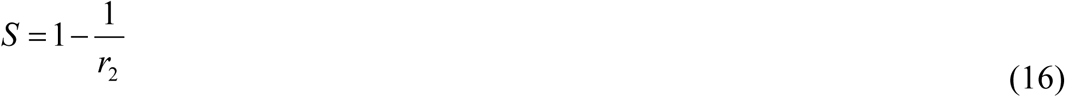

If only *P* and *N* are present, *N* always outcompetes *P*. If only *N* and *S* are present, *S* always outcompetes *N*. If only *P* and *S* are present, there is a particular density, *P = P_0_*, where the growth rates of *P* and *S* are equal:
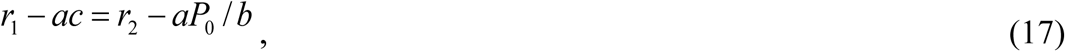

from which
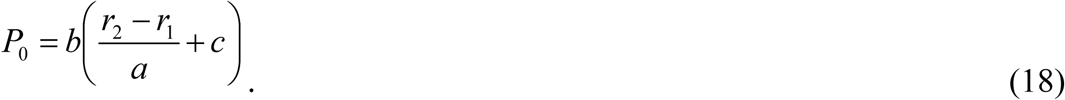

The corresponding density *S* = *S*_0_ at this point is
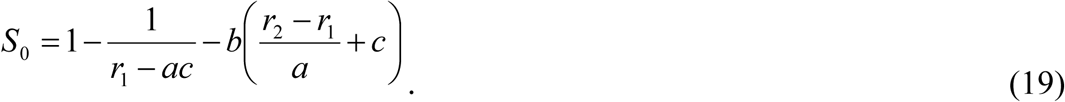

The point (*P_0_, S_0_*) is an unstable fixed point. The system moves away from this point to the stable point for either *P* or *S* alone. The unstable fixed point only exists if *S_0_* > 0 (in equation 19). This condition is true for values of *a* between *a_min_* and *a_max_*, where *a_min_* and *a_max_* can be determined by setting *S_0_* = 0 in equation 19. When *a* approaches either of these two limits, the fixed point for *P* becomes unstable. In summary, when considering the two cell types *P* and *S*, the fixed point for *S* (equation 16) exists for all *a* and is independent of *a*. The fixed point for *P* (equation 14) exists for *a_min_* < *a* < *a_max_* and is a decreasing function of *a* in this range. The reason that *P* can only outcompete *S* in the intermediate range is that if *a* is too low, the there is too little antibiotic to inhibit the growth of *S*, and if *a* is too high, the metabolic cost to *P* outweighs the benefit of inhibiting *S*. Note that in the intermediate range, the *P* solution will only arise if there is sufficient *P* present initially. A small amount of *P* cannot invade a population of *S* in the well-mixed case.

In the case where all three populations *P*, *S* and *N*, are present initially, the fixed point with only *S* (equation 16) is the only fixed point that remains stable. The only-*P* point is unstable to invasion by *N*. The only-*N* point is unstable to invasion by *S*. All trajectories thus end up at the only-*S* point. The conclusion is that antibiotic production should not be evolutionarily stable according to this well-mixed model because the producers are always invaded by non-producing cheats, which are then outcompeted by the sensitive species. Understanding the evolution of antibiotic production therefore requires us to account for spatial interactions. The spatial version of this model will be described in the following section.

## Description of the Stochastic Spatial Model

We consider a square lattice (of size L=1024×1024) in which each site can either be occupied by a cell or vacant. Cells may be either *P, N* or *S*. The growth rates of these cell types are *r_P_* = *r_1_* - *ac*, *r_N_* = *r_1_* and *r_S_* = *r_2_ - A_loc_*, where *A_loc_* is the local concentration of antibiotic on the site. The local neighbourhood of a site consists of that site plus the eight neighbouring sites in the Moore neighbourhood. The local concentration at any site is therefore *A_loc_* = *an_P_*/9*b*, where *n_p_* is the number of producers in the local neighbourhood. In some cases, a resistant cell type, *R*, that is not affected by the antibiotic, is also included. Its growth rate is *r_R_* = *r_2_* - *c_R_*, where *c_R_* is the cost of resistance. Later in the paper, we will include the possibility of mutation between *P* and *N* types, and between *S* and *R* types, but mutations from *P/N* to *R/S* are not considered possible.

The model proceeds in time steps, *δt*. In each time step, each site on the lattice is visited once in a random order. If there is a cell on the site, it reproduces with probability *r_i_δt*, where *i* is the type of cell being considered. If reproduction occurs, a site is chosen at random from the eight surrounding neighbours. If the neighbour site is vacant, the new individual is placed in this site. If the neighbour site is already occupied, it remains occupied by its current cell type, and the new individual is discarded.

We set the death rate of cells to be *v* = 1 for all cell types. Therefore, 1 time unit is the mean lifetime of a cell. In each time step, each cell has a probability *vδt* to die, giving rise to a vacant site. Patches of cells spread and move across the lattice as a result of birth and death events. We do not consider motion of individual cells independently of birth and death.

## The effect of antibiotic production rate and production cost on competition between *P*, *N* and *S* cell types

Simulations were performed on a lattice of 1024×1024 sites with *δt* = 0.01. Initially, sites were set randomly to be *P, N* or *S* with probability 5% each, and vacant with probability 85%. No *R* cells were present, and no mutation was possible. Simulations were run until only one cell type was present, or until a time *t* = 10,000 time units. Simulations were performed over a wide range of production rate *a*, and production cost *c*, with the other parameters being fixed at *r_1_* = 2, *r_2_* = 2.5, *v* = 1, and *b* = 10. Fig 1 shows several typical outcomes for different combinations of *a* and *c*, including cases where either *S* or *P* or *N* is the sole remaining cell type, and a case where all three remain present for a long time. There is a wide range of parameter values when all three types coexist. This occurs due to the formation of a spatial structure consisting of patches that are much larger than the individual lattice site – see Fig 2. On a local scale, there is cyclic dominance, where *P* replaces *S*, *S* replaces *N* and *N* replaces *P*. Hence, the patches move in space and time, but the overall system remains stable with three cell types.

**Fig 1.**
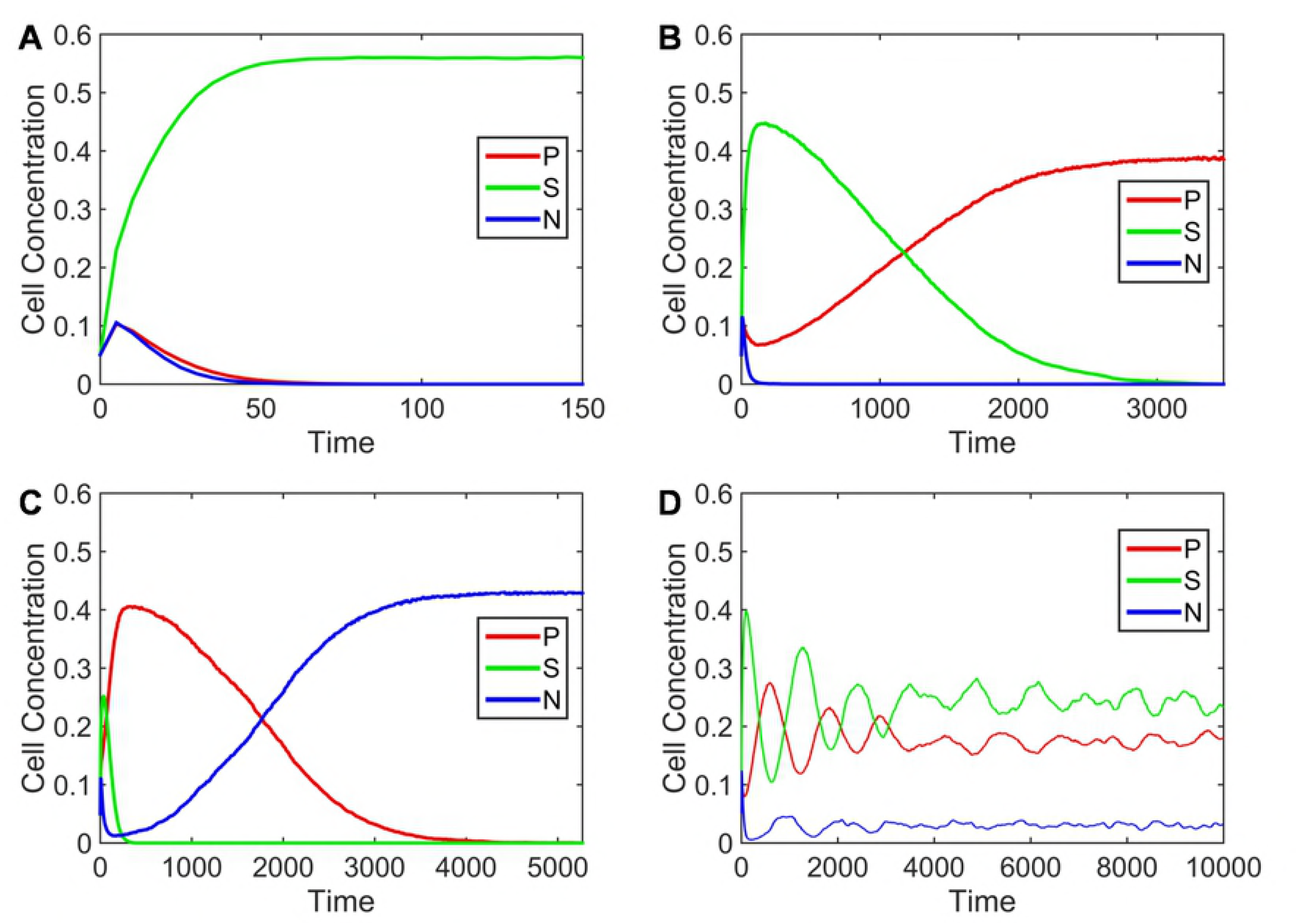
Cell concentrations as a function of time in different runs of the lattice model. Each run begins with randomly positioned *P, N* and *S* cells with a concentration of 0.05 each. For all examples *r_1_* = 2, *r_2_* = 2.5, *b* = 10 and the lattice size is 1024×1024. The other parameters differ. A) Sensitive cells win when *c* = 0.0011 and *a* = 30. B) Producers win when *c* = 0.0011 and *a* = 100. C) Non-producers win when *c* = 0.0002 and *a* = 90. D) The three cell types coexist when *c* = 0.001 and *a* = 150.

**Fig 2.**
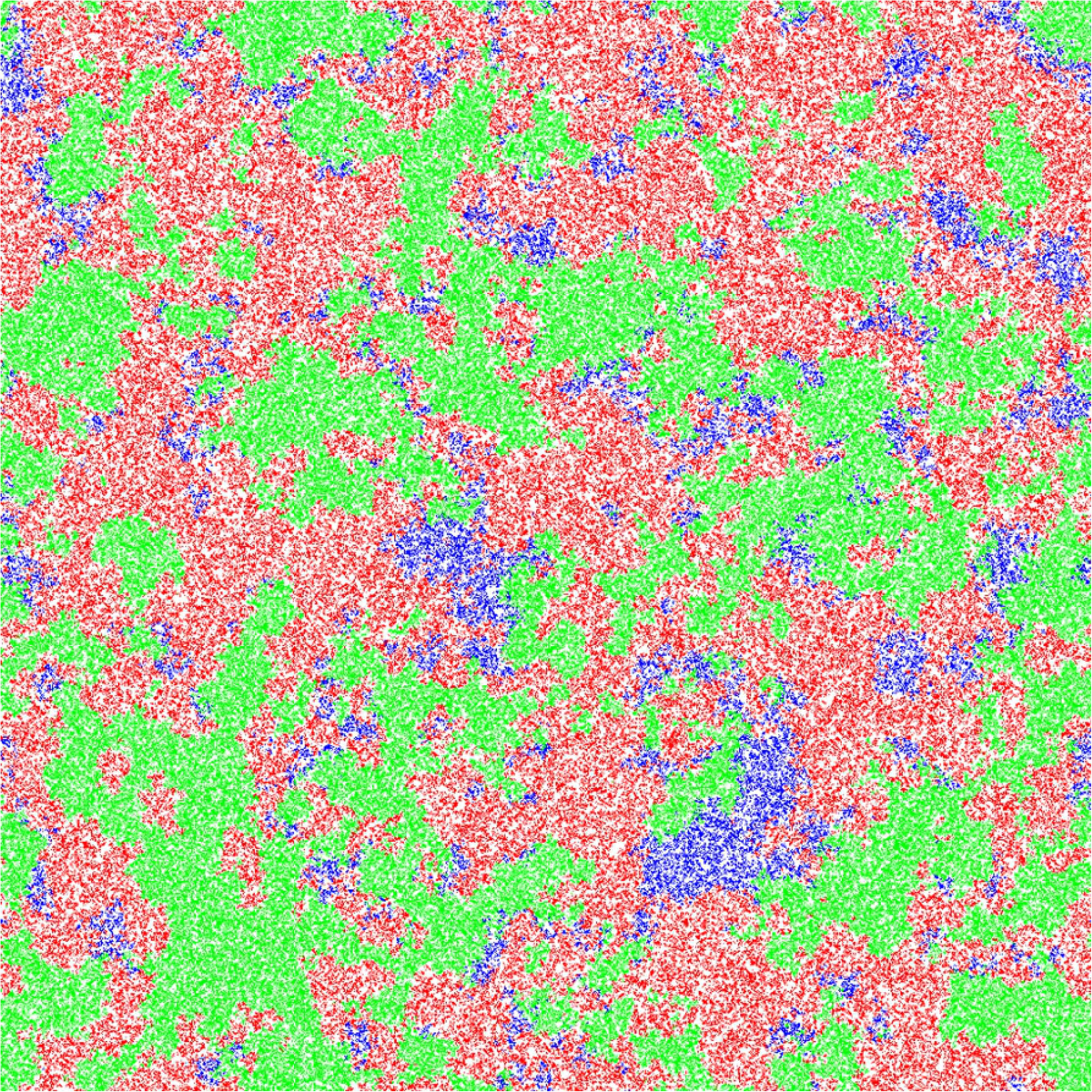
Snapshot showing the spatial structure of the lattice in a case where the three cell types coexist. The snapshot corresponds to the state of the system at time t = 4800 in Fig 1D. Parameters: *c* = 0.001; *a* = 150; *b* = 10; lattice size: L= 1024×1024.

There were some parameter combinations where the outcome was not always the same if the simulation was repeated. There is a tendency for the concentrations to oscillate, and sometimes the concentration of one cell type passes close to zero, especially at the beginning of the simulations before the spatial structure that stabilizes the coexistence is established. If one cell type dies, then one of the remaining cell types out-competes the other. Thus, the outcome can vary from run to run, even though the lattice size is quite large (1024×1024). To investigate the range of possible outcomes over the *a-c* parameter space, simulations were repeated 100 times for each *a-c* combination. The most frequent outcome from these trials, for each parameter set, is depicted by different colours in Fig 3.

**Fig 3.**
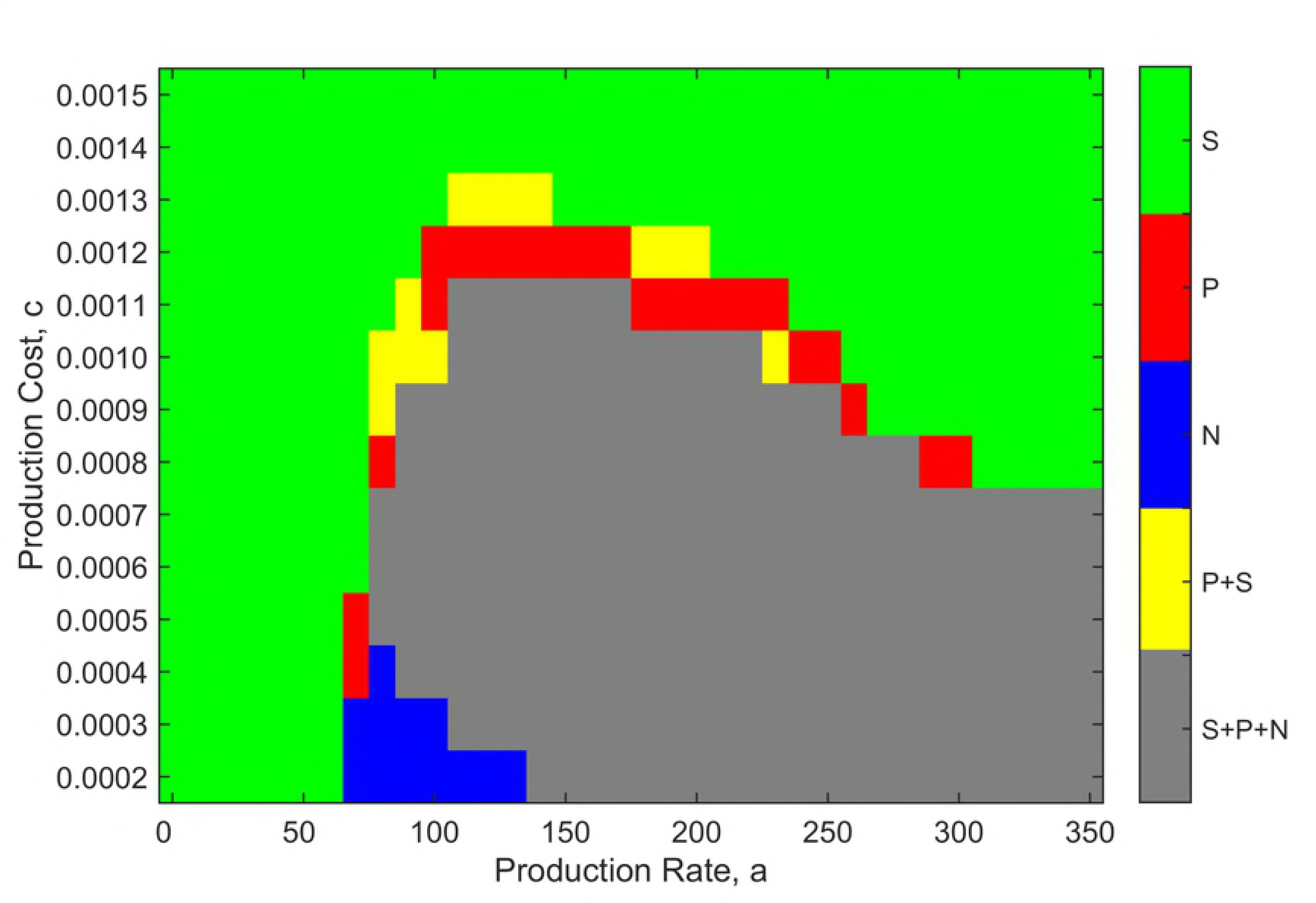
Majority outcomes of 100 runs of time 10,000 as a function of production rate *a* and production cost *c*. Green – *S* only; red – *P* only; blue – *N* only; grey – *S, P* and *N*; yellow – *S* and *P*. Parameters: *r_1_* = 2, *r_2_* = 2.5, *v* = 1*, b* = 10, L= 1024×1024, *δt* = 0.01.

From Fig 3, it can be seen that if the production rate is too low, sensitive cells win for all values of *c* because they grow faster than producers and there is insufficient antibiotic to reduce their growth rate. If the cost of production is too high, then sensitive cells win at all values of *a*, because the high cost of antibiotic production reduces the growth rate of the producers to a greater extent than the reduction in growth of S by the antibiotic. It can also be seen that for *c* ≥ 8 x 10^−4^, there is a value of *a* above which *S* always wins. We know that too high a production rate is always detrimental for producers, because eventually the growth rate will fall below the death rate, in which case the producer would not survive even if there were no competition. Thus, there must be a value of *a* above which *S* always wins. For costs less than *c* = 8 x 10^−4^, this value is beyond the range of *a* that we were able to simulate.

As long as the cost is not too high, there is an intermediate range of *a* where the three cell types coexist (depicted in grey). Close to the boundaries of this coexistence region, there are parameters for which the most frequent outcome is only *P* or only *N*. There are also cases (yellow) where *S* and *P* survive till the end of the simulation. This suggests that the fitnesses of these two cell types are close, and that the time required for one to out-compete the other is very long. Nevertheless, we expect that the outcome will always be either one cell type or three, after sufficient time.

Moving across Fig 3, for an intermediate value of cost *c* = 0.001, there are three principal regions where the outcome is predictable: *S* wins at low *a*, three cell types coexist at moderate *a*, and *S* wins at high *a*. The outcome is less predictable on the boundaries of the coexistence region. Fig 4 shows the frequencies of the different outcomes obtained from 100 trials for each value of *a* (with *c* fixed at 0.001). At both ends of the range, *S* wins in 100% of the runs (indicated by green bars), and in the central region, coexistence of all three cell types is observed in 100% of the trials (grey bars). The coexistence region is flanked by regions of variable outcomes.

**Fig 4.**
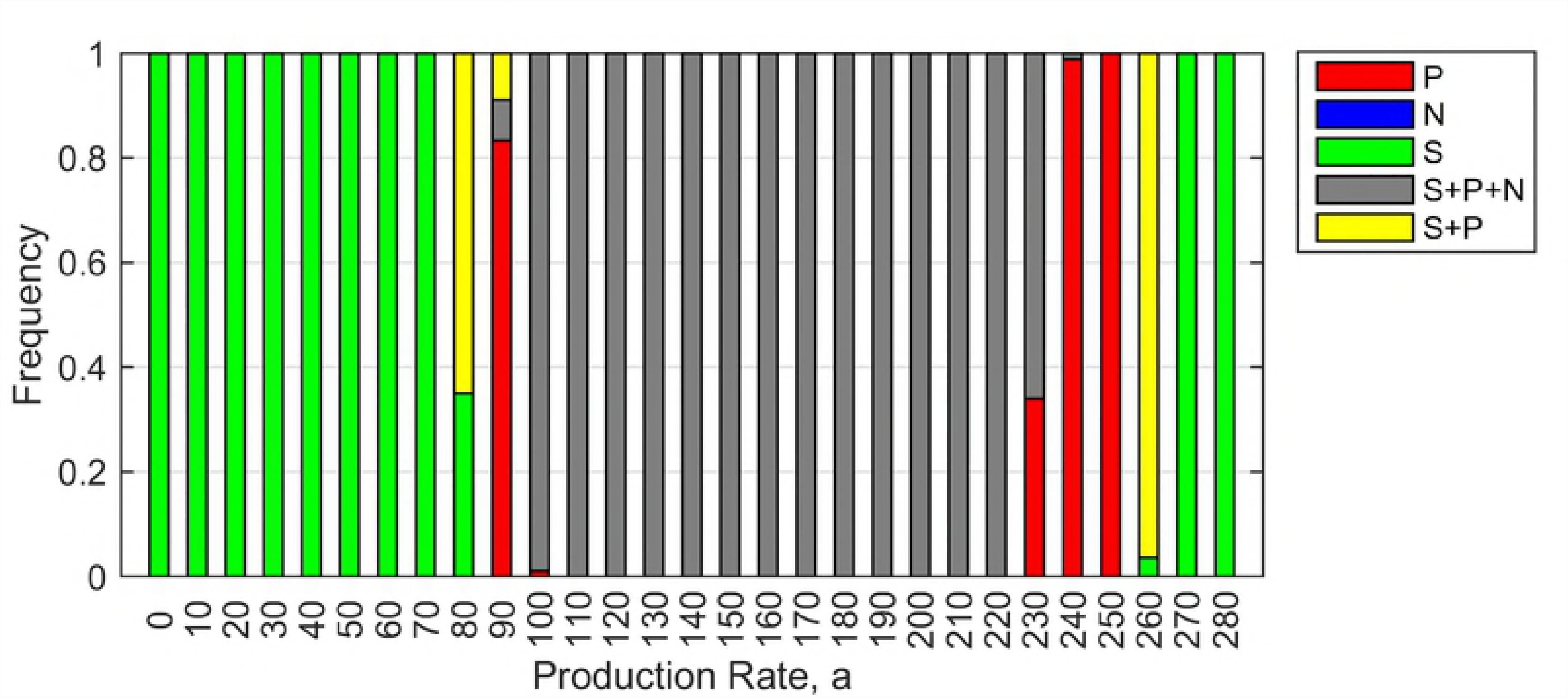
Possible outcomes of 100 simulation trials for each value of production rate *a*. Green – *S* only; red – *P* only; grey – *S, P* and *N*; yellow – *S* and *P*. Parameters: *r_1_* = 2, *r_2_* = 2.5, *v* = 1, *b* = 10, *c* = 0.001, *δt* = 0.01.

Fig 5 shows the time averaged concentrations for the cell types that survive at the end of the simulations. In the outer regions, where *S* wins, the *S* concentration is high and is independent of *a* (because there are no *P* cells). In the flanking regions the concentration of *P* is shown in the runs where *P* is the only surviving cell type. In the central coexistence region, as *a* increases, there is a greater advantage of *N* with respect to *P*. The ratio of *N:P* increases and the total fraction *P* + *N* decreases. The value of the production rate that maximizes the producer concentration is the minimal value that permits coexistence. If the production rate is higher than this, it benefits the sensitive cells not the producers.

**Fig 5.**
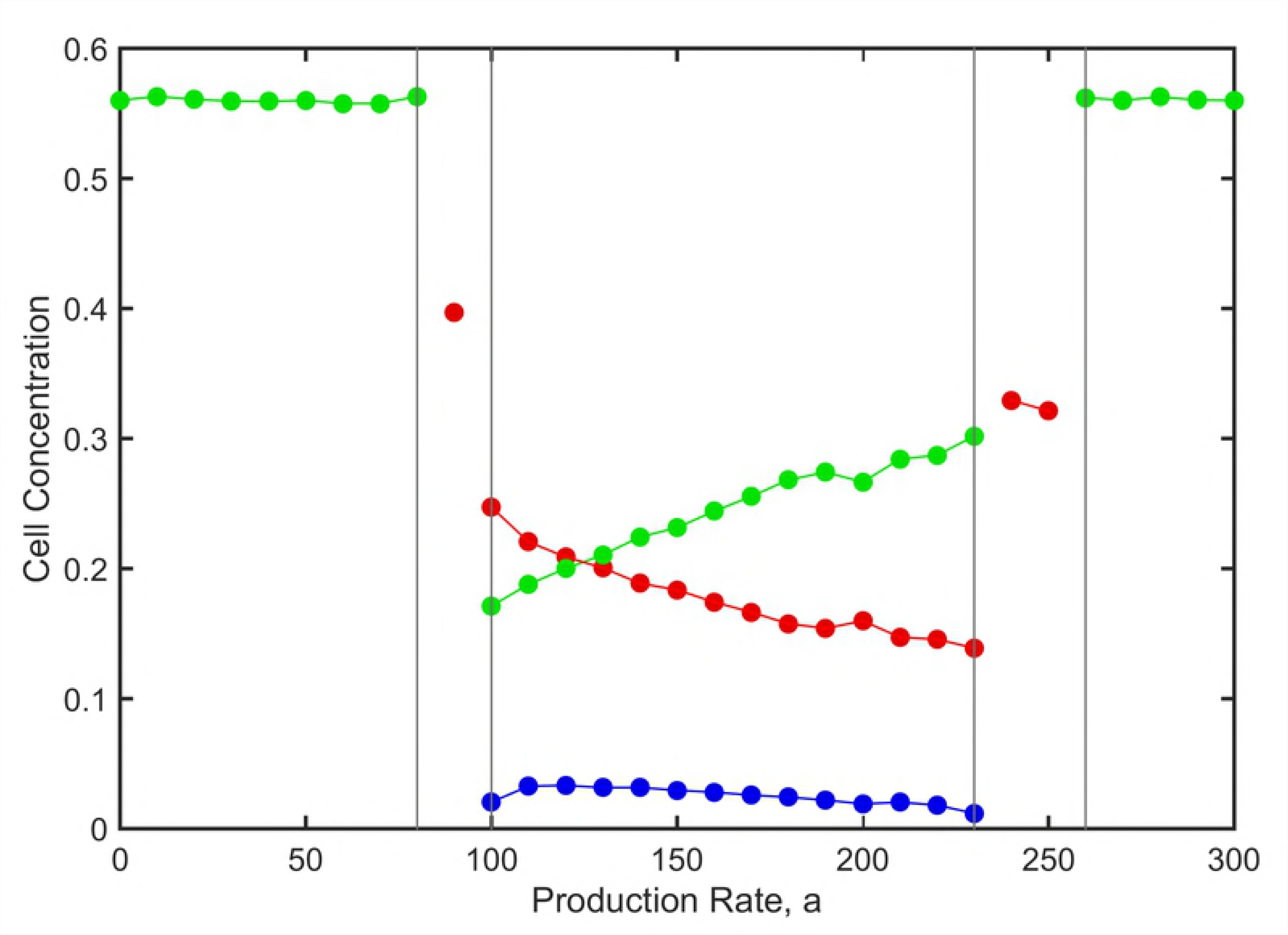
Time averaged cell concentrations for the surviving cell types as a function of *a*. Parameters as in Fig 4. P - red; N - blue; S - green.

## The effect of mutation between *P* and *N*

Non-producers are likely to arise frequently within a producer population due to mutations that inactivate the pathway of antibiotic synthesis or secretion. We therefore added the possibility of mutation between *P* and *N* cells to the model in the previous section. For simplicity, mutation was assumed to be equally likely from *P* to *N* and from *N* to *P*. When either a *P* or an *N* cell reproduced, the offspring cell was changed to the other cell type with probability *u* = 10^−4^. No mutation was possible between *S* and the other cell types because these are different species.

Figs 6 and 7, with mutation present, are comparable to Figs 4 and 5, with no mutation. The presence of mutation simplifies the outcomes. It is no longer possible for either *P* or *N* to exist on its own. The only possible outcomes are *S* alone (which occurs either for high *a* or low *a*), or coexistence of all three types (which occurs for intermediate *a*). The flanking regions with unpredictable outcomes are much less prominent in this case.

**Fig 6.**
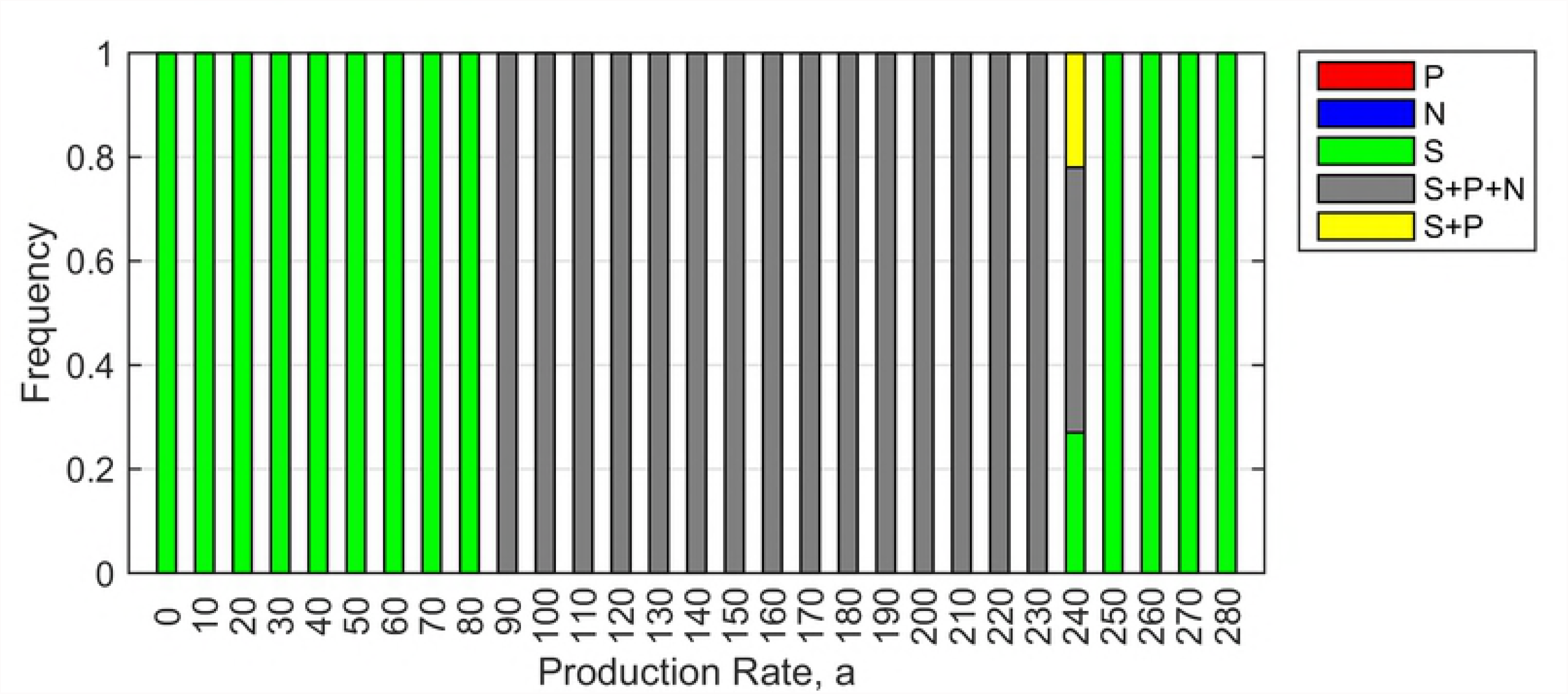
Possible outcomes of 100 simulation trials for each value of production rate *a* in the presence of mutation between *P* and *N*. Green – *S* only; red – *P* only; grey – *S, P* and *N*; yellow – *S* and *P*. Parameters: *r_1_* = 2, *r_2_* = 2.5, *v* = 1, *b* = 10, *c* = 0.001, *δt* = 0.01, µ=10^−4^.

**Fig 7.**
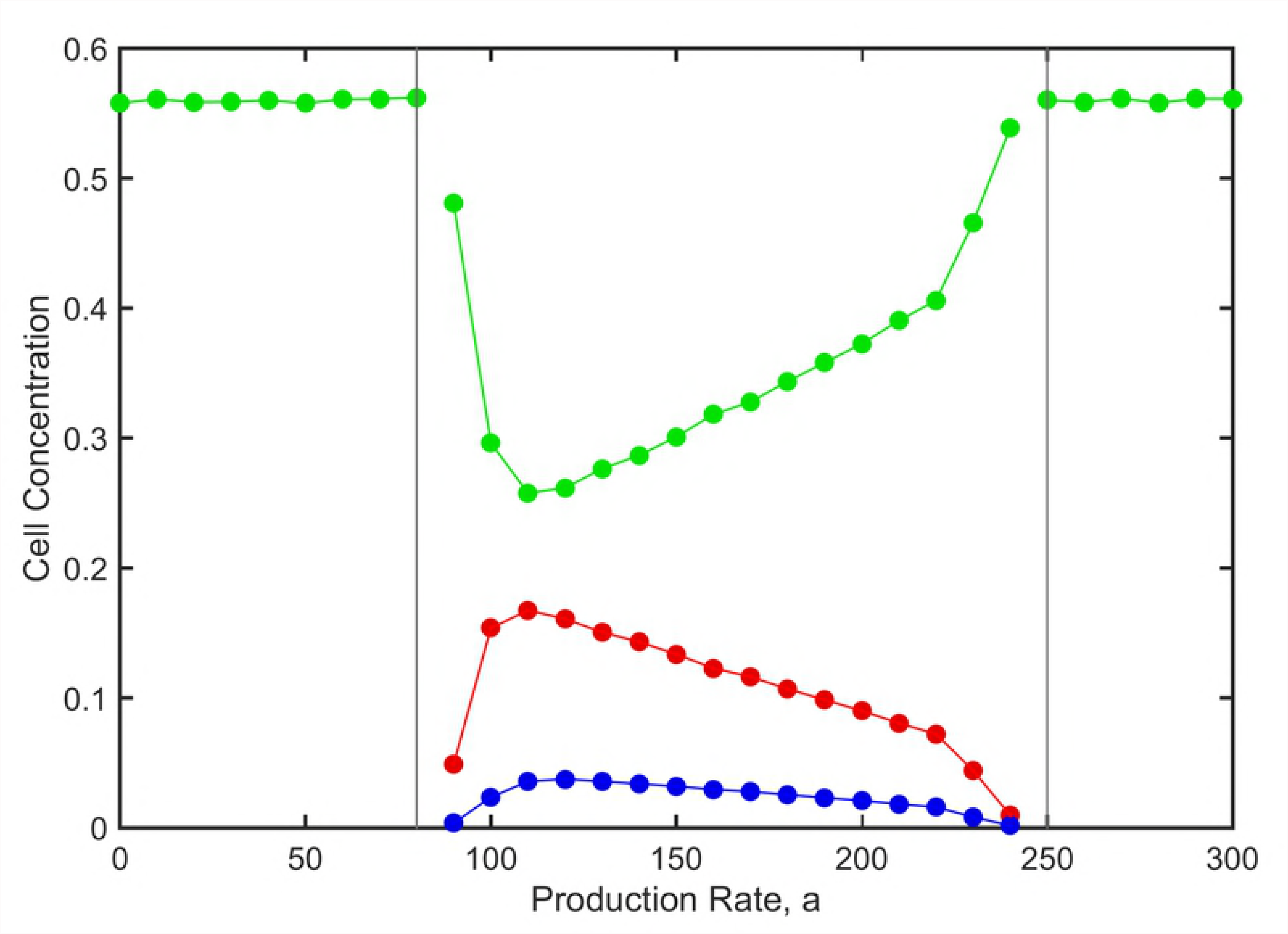
Time averaged cell concentrations for the surviving cell types as a function of *a* in the presence of mutation between *P* and *N*. Parameters as in Fig 6. P - red; N - blue; S - green

## Evolution of the rate of antibiotic production

In this section, we consider the evolution of the rate of antibiotic production *a* and the competition between strains of producers with different *a*. Fig 8 shows an example starting with producers of rate *a* = 110, non-producers (*a* = 0), and sensitive cells. This is a combination in which coexistence occurs and producers have a relatively high concentration (as shown in Figs 4–7). After 5,000 time units we allowed producers to mutate between their initial production rate and a production rate of *a* = 50. After a short time, the non-producers went extinct, and the two producers remained in coexistence with the sensitive cells (as shown in Fig 8).

**Fig 8.**
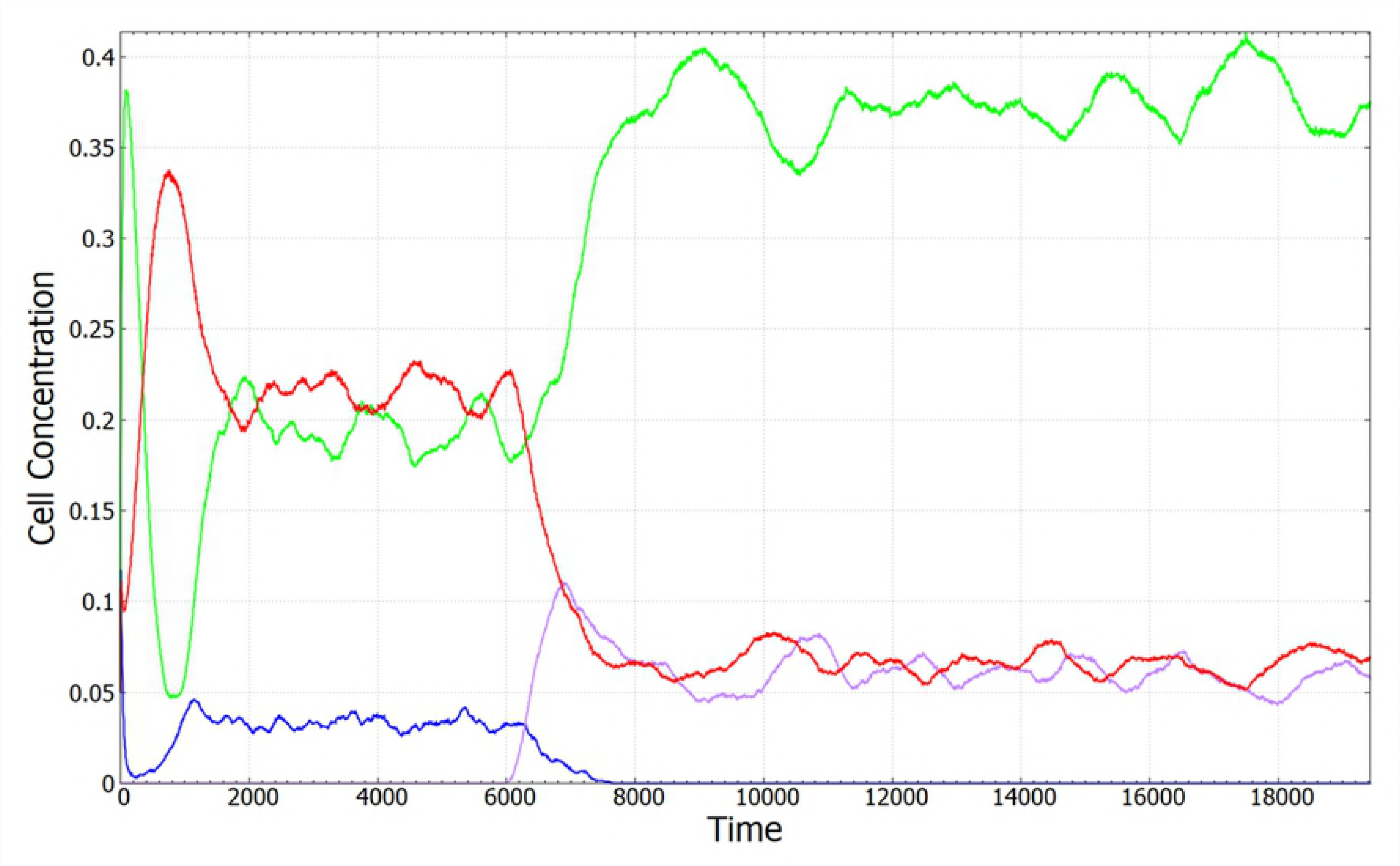
Time evolution of cell concentrations in a case where a low-rate producer replaces a non-producer. There are three cell-types initially: *P* with *a* = 110 (red)*, N* (blue), and *S* (green). At *t* = 6000, a low-rate producer with *a* = 50 (violet) is introduced. The low-rate producer replaces the non-producer and a new stable coexistence is established. Parameters: *b* = 10, *c* = 0.001, *δt* = 0.01, μ = 10^−4^, lattice size: L = 1024×1024.

We then considered simulations in which multiple types of producers are possible having all values of *a* in multiples of 10 between 0 and 200. We denote the producer types as *P_0_*, *P_10_*, *P_20_* …*P_200_*, where *P_0_* is equivalent to *N*. We considered three initial conditions in which the following cell types were present: (i) *P_150_*, *P_0_* and *S*; (ii) *P_200_*, *P_0_* and *S*; (i) all types from *P_0_* to *P_200_* and *S*. Simulations ran for 1000 time-steps without mutation, in order to allow the establishment of spatial structure. Mutation was then turned on among the producers with rate μ = 10^−4^. When a mutation occurred, the value of *a* was replaced randomly by one of the other possible values of *a* between 0 and 200. No mutation occurred between *S* and the *P*. In each case, these simulations evolved towards a stable state of coexistence of sensitive cells with multiple strains of producer. The time-averaged concentrations of the producer types are shown in Fig 9. The three initial conditions converge to similar end points. The most frequent producers are *P_110_* and *P_100_*, close to the values of *a* for which single producer strains do best in Figs 5 and 7. Producers with too high production rate do badly, since there is a growth rate penalty proportional to *a*, and there is no benefit to producing more antibiotic than necessary. There is a significant broad spread of low-rate producers across the range from *P_0_* to *P_90_*. These are cheats that can only survive due to the presence of *P_100_* and *P_110_*. This shows that evolution favours the spread of low-rate and zero-rate producers, but these cannot take over the population, and the mixture of low and high rate producers remains stable in the presence of the sensitive cells.

**Fig 9.**
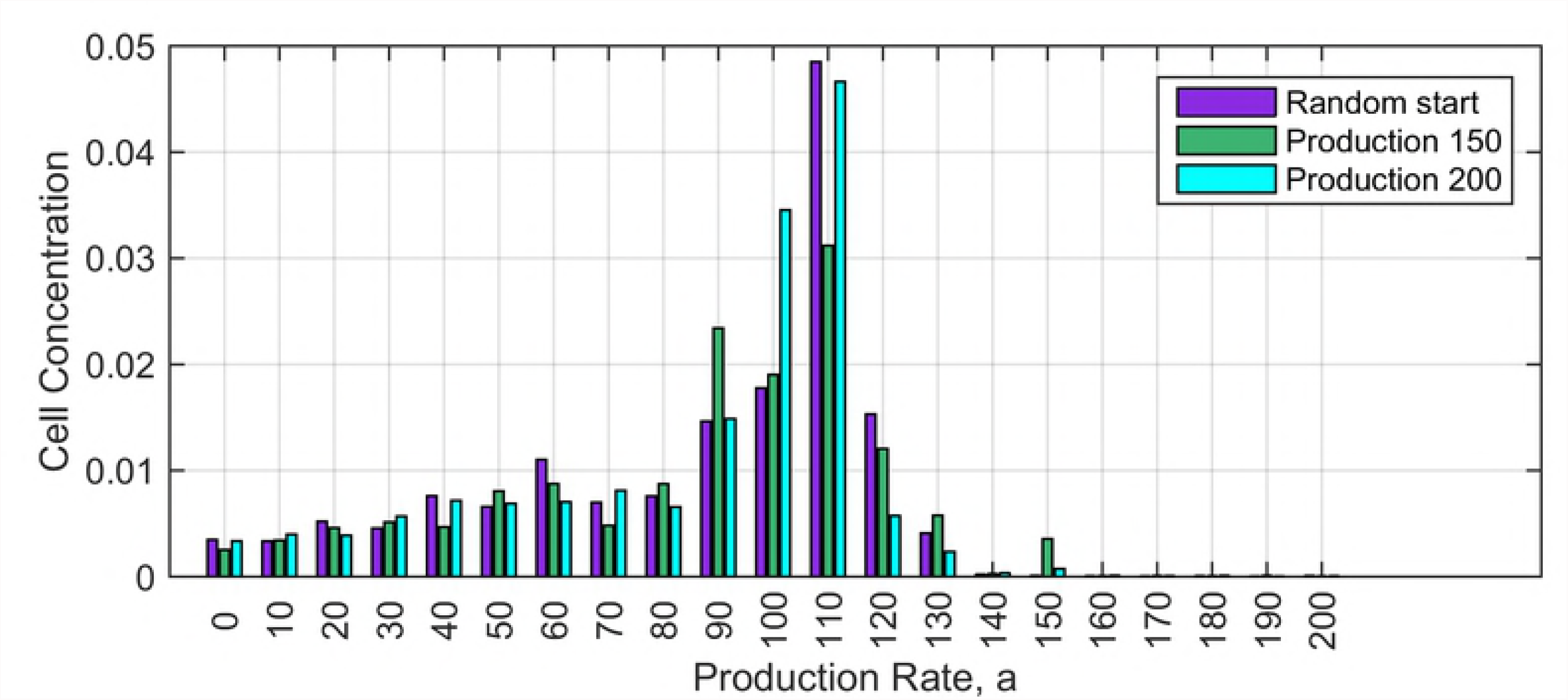
Time averaged cell concentrations in simulations with mutations between producers of many different production rates. Three sets of initial conditions are described in the text. These converge to similar distributions in the long-time limit. Simulations were run for 100,000 time units and an average was taken over the second half of the simulation. Parameters: *b* = 10, *c* = 0.001, *δt* = 0.01, μ = 0.0001, lattice: L = 1024×1024. The equilibrium concentration of *S* in these runs is close to 0.32.

## Introduction of a Resistant Cell Type

In this section, we consider a resistant cell type, *R*, that belongs to species 2. It has a growth rate *r_R_* = *r_2_* - *c_R_*, independent of the antibiotic concentration, where *c_R_* is the cost of resistance (relative to the sensitive cells). In absence of mutation, the resistant cell type, *R*, is similar to a non-producer, *N*, because both have a growth rate between *P* and *S*. Thus, in absence of mutation, coexistence of the three types *P, R* and *S* can occur in a similar way to the three types *P, N* and *S*. Our objective here is to consider the case with four possible cell types *P, N, R* and *S* in the presence of mutations. In this case, it is significant that *N* evolved from a producer strain, so that mutations occur between *P* and *N*, but *R* evolved from a sensitive strain, so that mutations occur between *S* and *R*. Thus, *R* and *N* are no longer equivalent due to their distinct evolutionary origins, and it makes sense to consider both in the same simulation. From the point of view of species 2, the sensitive cells are cheaters. If resistant cells are sufficiently common, the producers will be kept to a low level, in which case there is an opening for sensitive cells that cheat by avoiding the cost of resistance.

We began with three cell types *P*, *N* and *S* and allowed the system to establish a spatial structure with coexistence. After 5000 time units, mutation was turned on between *P* and *N*, and between *S* and *R*, with mutation rate μ = 1 x 10^−4^. We then determined whether the *R* type could invade the system and cause it to shift to a new steady state. As before, we set *r*_*N*=_ *r_1_* = 2.0 and *r_S_* = *r_2_* = 2.5. With *a* = 110 and *c* = 0.001, the growth rate of the producers is *r_P_* = 1.89. We considered separate runs of the simulation with growth rates of the resistant cells in the range *r_R_* = 1.5 - 2.5, which corresponds to costs of resistance between 1.0 and 0.0.

For *r_R_* < *r_P_*, the resistant strain has the lowest growth rate. There is a near-zero equilibrium concentration of *R*, because resistant mutations cannot spread. For *R* to spread, it is necessary to have *r_R_* > *r_N_*, not just *r_R_* > *r_P_*. At *r_R_* = 1.9, we have *r_P_* < *r_R_* < *r_N_*. If there were no non-producer, the resistant strain could spread at this point. However, in the presence of non-producers, the resistant strain still cannot spread. For the data point at *r_R_* = 2.0, *R* and *N* have equal growth rates. Nevertheless, the mean concentration of *R* is still very low and much less than *N*. This is probably due to a spatial effect. As *R* cells are created by mutations from *S*, they usually appear in the middle of patches of *S*, where they cannot spread because *r_R_* < *r_S_*. On the other hand, *N* cells usually arise in the middle of patches of *P*, from where they can immediately spread due to their selective advantage relative to *P*.

When *r_R_* > *r_N_* the concentration of *R* cells becomes significant, but still remains low compared to *S* and *P* for the case shown in Fig 10. As *R* now outgrows both *P* and *N*, its presence has a significant negative effect on both these strains. Both *P* and *N* decrease in concentration as *r_R_* increases above 2.0. However, the reduction in the number of producers makes it easier for sensitives. Thus, the chief effect of increasing the growth rate of *R* is to increase *S*. There comes a point (close to *r_R_* = 2.3 in this example) where *R* causes the extinction of *P* and *N*. At this point, there is no benefit to resistance; therefore, the whole system is taken over by *S*. Finally, when *r_R_* = *r_S_*, there is no cost of resistance. Therefore, *R* and *S* are indistinguishable, and either can take over the population with a probability proportional to their frequency as expected in a neutral evolution scenario.

**Fig 10.**
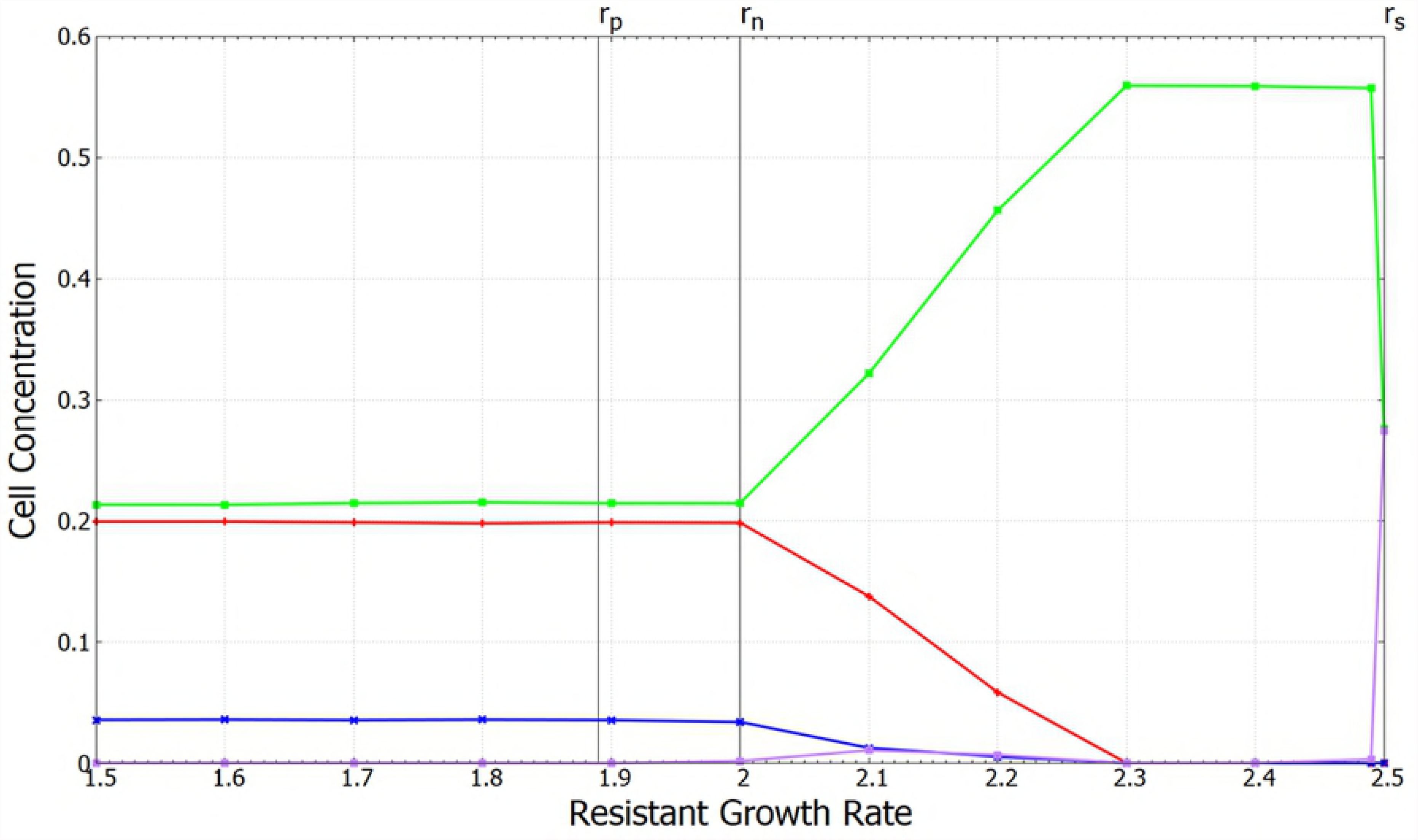
Equilibrium concentrations of four cell types *P* (red), *N* (blue), *S* (green) and *R* (violet). Mutations occur between *P* and *N*, and between *S* and *R*. Parameters: *r_1_* = 2.0, *r_2_* = 2.5, *a* = 110 *b* = 10, *c* = 0.001, *δt* = 0.01, μ = 10^−4^, and lattice size: L= 1024×1024.

In summary, resistance mutations can persist only if the growth rate of the resistant strain is greater than the non-producing strain of the producer species, and not simply greater than the producing strain. Coexistence of all four cell types is possible for a limited range of resistant cell growth rates. In this range, the presence of the resistant strain has a significant effect on the balance between the other strains, even though the resistant cells remain fairly rare. Coexistence is not possible if the resistant cells grow too rapidly, because in this case, they cause extinction of the producers, and sensitive cells take over the whole system.

## Discussion

Drawing on previous models of colicin and rock-paper-scissors systems, we have created a spatial model that allows for the study of interaction between producer, non-producer, sensitive and resistant cells. The three cell types *P, N* and *S* coexist for an intermediate range of antibiotic production rate. If the production rate is either too high or too low, only the sensitive cells survive. Coexistence of producers and sensitive cells is only possible if the non-producers are also present because the system with only two cell types would be taken over by either *P* or *S* depending on the production rate of the antibiotic. By allowing multiple strains of producers to evolve over a broad range of production rate, we have shown that producers with a moderately large production rate can survive against cheater mutations that immediately reduce production rate to zero *and* against mutations that gradually reduce production rate in small amounts. In the latter case, we have shown that the stable state is likely to consist of a mixture of strains with a principle producer of relatively high rate and many strains with low or zero production rate that survive alongside the principle producer but do not eliminate it.

We have also shown that for certain parameter ranges, the four cell types *P, N, R* and *S*, may all coexist. This may be thought of as an example of the evolution of cooperation in two senses. Maintenance of the antibiotic production mechanism in species 1 requires that producers (cooperators) survive the competition of non-producers (cheats). Maintenance of a resistance gene in species 2 requires that the resistant cells (cooperators) survive the competition of sensitive cells (cheaters). Furthermore, both mechanisms can only be maintained when both species survive. If species 1 dies out, there is no benefit to resistance, and species 2 will become entirely sensitive. If species 2 dies out, there is no benefit to antibiotic production, and species 1 will become entirely non-producing.

A large variety of both antibiotic production mechanisms and resistance mechanisms is found in nature. The model we studied here identifies the minimal requirements for the sustenance of both producer and resistant phenotypes in a bacteriostatic model of microbial interactions - namely, a spatially distributed system in which there are local interactions between nearby cells. This allows for a patchy structure of cell types to emerge in which there can be cyclic dominance of three (or more) cell types. In a well-mixed model, neither resistance nor production phenotypes would be stable.

There are several more complex factors not included in our model that are likely to influence the survival of production and resistance mechanisms in the real world. The antibiotic molecule may diffuse over a range considerably wider than the cell size. The release of antibiotic into the environment may clear a wide space around the producer colony. Additionally, if the antibiotic had a long life-time in the environment, this would prevent colonisation by sensitive species of a region previously occupied by producer strains. The results presented here use a model where producers affect sensitive cells on neighbouring lattice sites only. One lattice site may be thought of as an effective range of the antibiotic (*i.e.* the typical distance that it diffuses before breaking down). We assumed that production and breakdown of antibiotic is rapid compared to cell division, so that the local antibiotic production is proportional to the local concentration of producers at the current time step. We have also studied a more complex model in which these assumptions are relaxed. In that model, diffusion of the antibiotic beyond nearest-neighbour sites is incorporated explicitly resulting in variable antibiotic concentration across the lattice. We keep track of the concentration of antibiotic on each lattice site since it affects the birth rates of *P* and *S*. We found that this model was much slower to simulate, because updating the antibiotic concentration on each site requires significant time, and because the spatial size of the patches becomes larger when the range of interaction becomes larger than nearest neighbour. This means that larger lattice sizes are required to get reproducible results, which also increases simulation time. We have obtained similar results to Figs 3 and 4 with the more complex model, but with fewer parameter combinations, and with a stronger influence of finite size effects. Therefore, we do not present these results here. Nevertheless, it is worth pointing out that incorporating diffusion explicitly in our model does not prevent multi-species coexistence.

A natural extension of our work would involve considering the effect of alternative mechanisms of antibiotic resistance on multi-species coexistence in the bacteriostatic model. An important factor for maintenance of resistance mechanisms is that they are often encoded by genes carried on plasmids. The horizontal transmission of resistance via plasmids would permit rapid spread of resistance in cases where sensitive cells experienced a sudden introduction of antibiotic, although in many cases where there is a stable level of antibiotic present, it would pay to transfer the resistance gene from a plasmid to the bacterial chromosome, because the cost of the resistance mechanism itself is likely to be much less than the cost of carrying the plasmid in the cell [24]. Resistance can also be conferred without horizontal gene transfer. For instance, certain microbes are capable of producing and releasing an enzyme that degrades the antibiotic [25,26]. The spillover effect of degradation of the antibiotic by such species benefits the sensitive strains (cheaters) that can grow uninhibited in the antibiotic-free zone created around such degraders. In such cases, the diffusion of the antibiotic as well as the degrader enzyme is likely to affect the evolutionary dynamics of the multi-species systems.

Finally, stability of the microbial ecosystem should depend on the manner in which individual species utilize and compete for nutrients. A recent study [27] has shown that species diversity can be maintained if multiple species can utilize the metabolic by-products of other coexisting strains as nutrients in addition to the resource available in the ecosystem. Competition for nutrients, the efficiency with which individual species utilize available resources as well as inter-species conflict and cooperation are all likely to have a significant impact on the diversity and stability of microbial ecosystems.

## Acknowledgements

We thank Jonathan Dushoff, Tyler Meadows, Jonathon Stone, Gail Wolkowicz, and Gerry Wright for useful discussions on this project, and Andrew Tupper for help with implementing parallel processing versions of the lattice models.

